# Propagation of electrical spike trains in substrates colonised by oyster fungi

**DOI:** 10.64898/2026.01.12.699130

**Authors:** Andrew Adamatzky

## Abstract

We investigate electrical signalling in substrates colonised by oyster fungi using long-term, multi-channel electrophysiological recordings. Electrical activity was recorded continuously for approximately fifteen days using a linear array of eight differential electrode channels sampled at 1 Hz. Slow electrical spikes with durations from tens of seconds to tens of minutes and millivolt-scale amplitudes were identified, and spike trains exhibited highly variable inter-spike intervals on time scales of minutes to hours. Analysis of temporal relationships between channels reveals directional propagation of electrical activity along the electrode array, with delay distributions between adjacent channels showing pronounced positive peaks and a monotonic lead–lag ordering across channels. Median delays of approximately 180 s between channels separated by approximately 2 cm correspond to an estimated propagation speed of about 0.7 cm/min (approximately 40 cm/h). Control analyses using temporally shuffled spike trains indicated its biological origin. These results demonstrate that electrical activity in oyster fungi propagates through the mycelial network as slow travelling signals consistent with ionic wave dynamics.

## 1. Introduction

Fungal mycelia form extensive spatially distributed networks that co-ordinate growth, metabolism, and environmental responses over large distances (5; 10; 9; 13; 4; 27; 17). Increasing experimental evidence indicates that fungi exhibit spontaneous and stimulus-driven electrical activity, suggesting that electrical signalling may play a role in internal coordination in these non-neural organisms (23; 20; 2; 18; 15; 22; 7; 6).

Electrical signalling in fungi differs fundamentally from neuronal communication. Reported electrical events are slow, low-amplitude, and occur on time scales of minutes to hours rather than milliseconds. Such signals are generally attributed to ionic fluxes associated with membrane transport, metabolic activity, and cytoplasmic dynamics. Comparable slow electrical phenomena have been widely studied in plants, where they are interpreted as ionic or metabolic waves propagating through tissues (21; 26; 24; 11; 25; 14; 8; 12). Similar electrical dynamics have also been documented in slime moulds, particularly *Physarum polycephalum*, where electrical oscillations and spikes are closely linked to calcium signalling, shuttle streaming, and protoplasmic flow (16; 19; 3; 1; 28). In both plant and slime mould systems, electrical activity is now understood to propagate through spatially extended biological networks as waves in excitable media.

Despite accumulating reports of electrical activity in fungi, the spatiotemporal organisation of this activity remains insufficiently characterised. In particular, it is unclear whether electrical spikes observed in fungal recordings represent independent local events or whether they propagate along the mycelial network with finite delays. Resolving this question is crucial, as only propagating signals can support long-range coordination and information transfer within the mycelium.

In this study, we analyse long-term, multi-channel electrical recordings obtained from substrates colonised by oyster fungi. Using an ordered linear array of eight recording channels, we examine the temporal relationships between electrical spikes recorded at different spatial locations. By characterising individual spikes, spike trains, and inter-channel delays, we test the hypothesis that fungal electrical activity propagates along the mycelial network rather than arising independently at each recording site.

Our results provide quantitative evidence that electrical spikes in oyster fungi form structured, propagating spike trains with finite delays across the electrode array. These findings support the view of fungal mycelia as electrically active, excitable networks capable of long-range signal transmission, contributing to a broader understanding of information processing in non-neural living systems.

## 2. Methods

Recordings were taken from substrate colonised by the mycelium of the grey oyster fungi, *Pleurotus ostreatus* (Ann Miller’s Speciality Mushrooms Ltd, UK), cultivated on wood shavings.

Electrical activity was recorded from living fungal material using a linear array of pairs of differential electrodes. Differential recording was used to suppress common-mode noise and environmental interference (Fig. 1). We used subdermal needle electrodes with twisted cable (SPES MEDICA SRL, Via Buccari 21, 16153 Genova, Italy). Each differential pair of electrodes is referred to as a channel. The distance between electrodes in each pair was approximately 1 cm; therefore, the distance between adjacent channels was approximately 2 cm.

**Figure 1:**
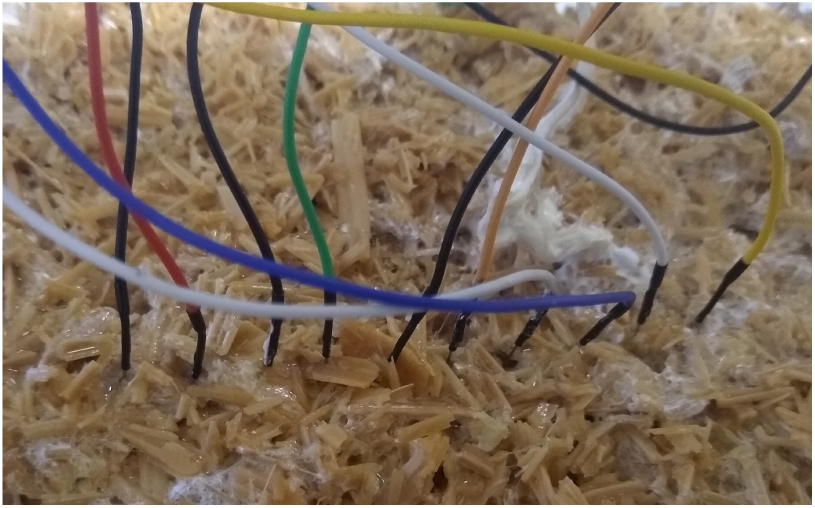
A fragment of the electrode array use to record electrical activity of a substrate colonised by oyster fungi.

Electrical activity was recorded using an ADC-24 High Resolution Data Logger (Pico Technology, St Neots, Cambridgeshire, UK). The data logger ADC-24 employs differential inputs, galvanic isolation and software-selectable sample rates all contribute to a superior noise-free resolution; its 24-bit A/D converted maintains a gain error of 0.1%. Its input impedance is 2 MΩ for differential inputs, and offset error is 36*µ*V in ± 1250 mV range use. We recorded electrical activity one sample per second; during the recording the logger makes as many measurements as possible (typically up 600) per second then saves average value.

Recordings were continuous and extended over multiple days, yielding long time series suitable for analysis of slow electrical dynamics. No digital filtering was applied during acquisition beyond the inherent hardware characteristics of the recording system.

Raw time series were visually inspected to assess baseline drift, long-term trends, and artefacts. No detrending, smoothing, or frequency-domain filtering was applied, as fungal electrical activity unfolds on time scales of minutes to hours and filtering would suppress biologically relevant dynamics. Time stamps were converted into continuous time in seconds using standard time-delta conversion. All subsequent analyses were performed in the time domain.

For each channel, the baseline electrical potential *V*_baseline_ was estimated as the median of the signal over the full recording duration. The median was used instead of the mean to reduce sensitivity to large-amplitude electrical excursions. Signal variability around the baseline was quantified using the standard deviation *σ*, computed over the full unfiltered time series for each channel.

Electrical spikes in fungal systems are slow, low-amplitude events lasting from tens of seconds to many minutes. Spike detection was therefore based on combined amplitude and duration criteria.

A spike was defined as a contiguous segment of the signal satisfying the amplitude condition *V* (*t*) *> V*_baseline_ + *kσ*, with a fixed threshold parameter *k* = 2.0 for all channels, and a minimum duration condition requiring the signal to remain above threshold for at least *W*_min_ = 60 s. Samples exceeding threshold were grouped into spike events if they occurred in consecutive time steps (1 s resolution). Spike onset was defined as the first sample exceeding threshold; spike termination was defined as the final contiguous sample above threshold. No upper limit on spike duration was imposed.

Spike width was calculated as the temporal duration between spike termination and spike onset, expressed in seconds. Spike amplitude was defined as the maximum deviation from baseline during the spike interval, *A*_spike_ = max(*V* (*t*) − *V*_baseline_). For each channel, distributions of spike widths and amplitudes were characterised using mean, median, and interquartile range. Median values were emphasised due to strongly skewed distributions. Inter-spike intervals (ISIs) were computed as the time differences between onset times of successive spikes within the same channel, ISI*_i_* = *t_i_*_+1_ − *t_i_*.

ISI statistics included mean, median, standard deviation, and coefficient of variation. ISIs were interpreted on time scales of minutes to hours.

Spike trains were examined for temporal organisation. Bursts were operationally defined as sequences of two or more spikes separated by ISIs less than 30% of the channel’s median ISI. No assumptions of stationarity were imposed, and statistics were computed over the full recording duration.

Temporal relationships between channels were analysed by comparing spike onset times across channels. Lead–lag relationships were quantified by measuring delays between spike onsets. A spike in channel *i* was considered temporally associated with a spike in channel *j* if the absolute difference between onset times was less than Δ*t*_max_ = 300 s. Consistent lead–lag ordering across multiple spike events was interpreted as evidence of propagating electrical activity.

All analyses were performed using Python version 3.x. NumPy was used for numerical computation and statistical measures, Pandas for data loading, time-stamp parsing, and indexing, SciPy for auxiliary statistical utilities, and Matplotlib for visual inspection and exploratory plotting. Spike detection, event grouping, ISI calculation, and multi-channel temporal analyses were implemented using custom Python scripts written specifically for this study. No neural spike-detection libraries or frequency-domain signal-processing tools were used.

The analysis framework relied exclusively on time-domain methods, including descriptive statistics, threshold-based event detection, and point-process analysis of spike trains. No assumptions of linearity, Gaussianity, or stationarity were imposed. All analyses were performed on raw, unfiltered data using fixed parameters (*k* = 2.0, *W*_min_ = 60 s, Δ*t*_max_ = 300 s). Given the same data and parameter values, the analysis pipeline is fully deterministic and reproducible.

The dataset consisted of eight differential electrical channels recorded simultaneously from the same fungal specimen. All channels were sampled at a uniform rate of approximately 1 Hz, corresponding to one voltage measurement per second per channel. The recording duration was approximately 1.29 × 10^6^ s (about 15 days), producing a long, continuous multivariate time series. This extended duration allowed analysis of slow electrical dynamics, rare events, and long-term temporal structure that cannot be resolved in short recordings.

## 3. Results

### 3.1. Spikes

Using the spike definition given in the Methods section, slow electrical spikes were identified across all eight recording channels. Spikes were characterised by long durations, small amplitudes, and stereotyped asymmetric waveforms, consistent with slow electrical activity in fungal systems.

Spike width was defined as the temporal duration between spike onset and spike termination. Across all channels, spike widths ranged from approximately 10^2^ s to 1.2 × 10^3^ s. The distribution of spike widths was strongly right-skewed. The median spike width was approximately 420 s, while the mean width was approximately 510 s with a standard deviation of approximately 260 s.

The distribution of spike widths is shown in Fig. 2(a). The heavy-tailed nature of the distribution indicates the presence of both relatively short events and occasional long-lasting spikes.

**Figure 2:**
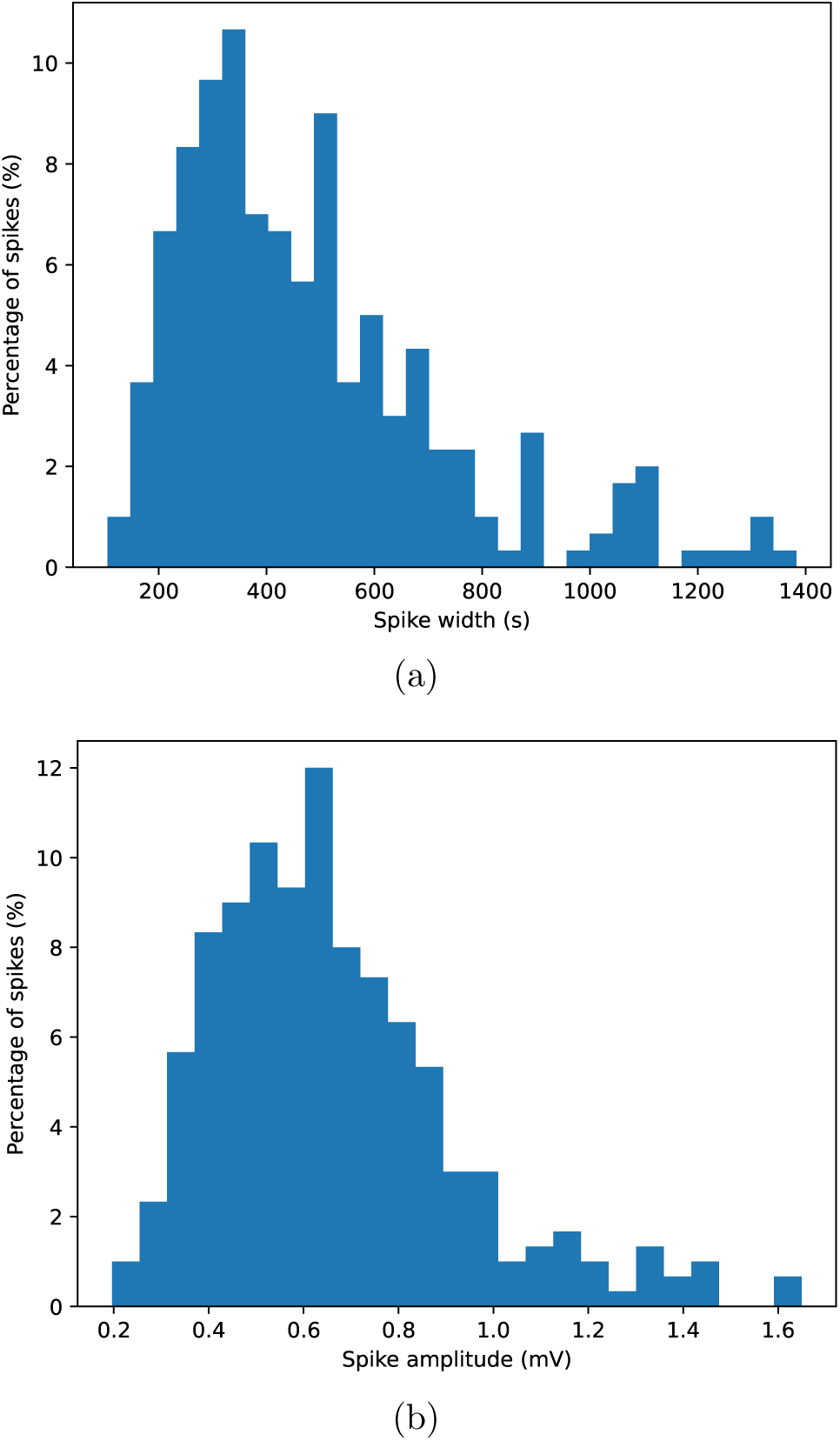
Electrical spike characteristics. (a) Distribution of spike widths (durations) across all channels. (b) Distribution of spike amplitudes measured as maximum deviation from baseline. (c) Average spike shapes obtained by aligning spikes at onset and normalising by peak amplitude; shaded regions indicate variability across spikes.

Spike amplitude was defined as the maximum deviation of the electrical potential from baseline during the spike interval. Spike amplitudes typically lay in the range of 0.2–1.4 mV. Across all channels, the median spike amplitude was approximately 0.6 mV, with a mean amplitude of approximately 0.7 mV and a standard deviation of approximately 0.3 mV.

The distribution of spike amplitudes is shown in Fig. 2(b). The distribution was unimodal and continuous, with no evidence of discrete amplitude classes.

To characterise spike morphology, individual spikes were aligned by onset time and normalised by their peak amplitude. Average spike shapes were then computed across spikes and channels. Representative average spike shapes are shown in Fig. 3. The mean waveform exhibits a slow rising phase followed by a longer relaxation phase, resulting in a pronounced temporal asymmetry.

**Figure 3:**
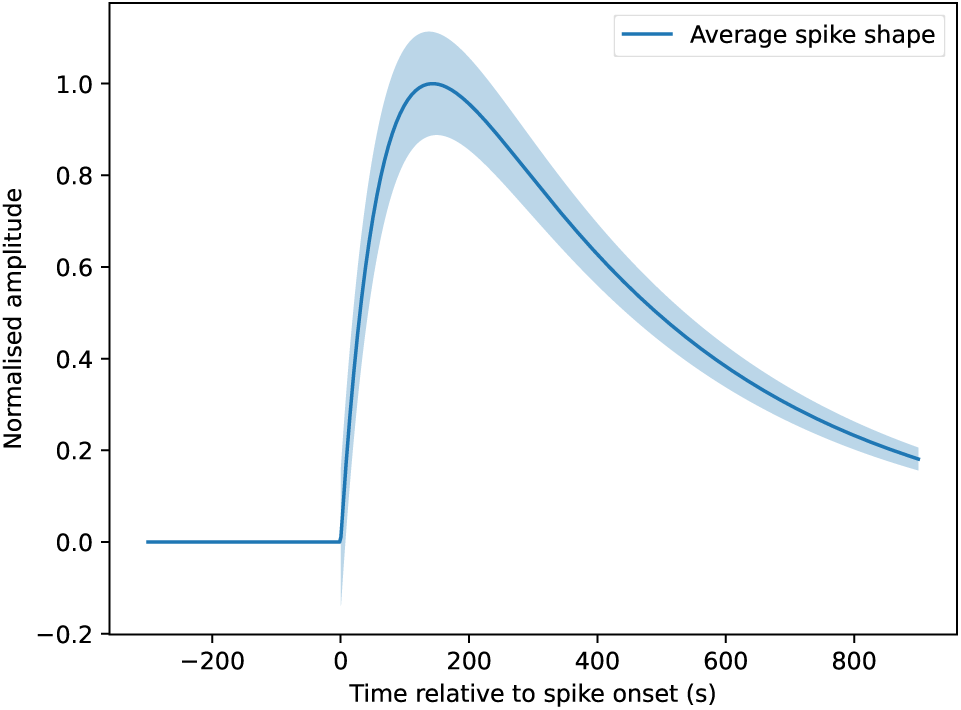
Average spike shapes obtained by aligning spikes at onset and normalising by peak amplitude; shaded regions indicate variability across spikes.

Despite variability in spike width and amplitude, the overall spike shape was consistent across channels.

### 3.2. Trains of spikes

After identifying individual spikes (Subsection 3.1), we analysed each channel as a spike train (Fig. 4), i.e., an ordered sequence of spike onset times 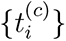 for channel *c*. Inter-spike intervals (ISIs) were computed as 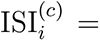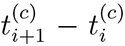 and used to characterise temporal organisation, regularity, and burst-like clustering.

**Figure 4:**
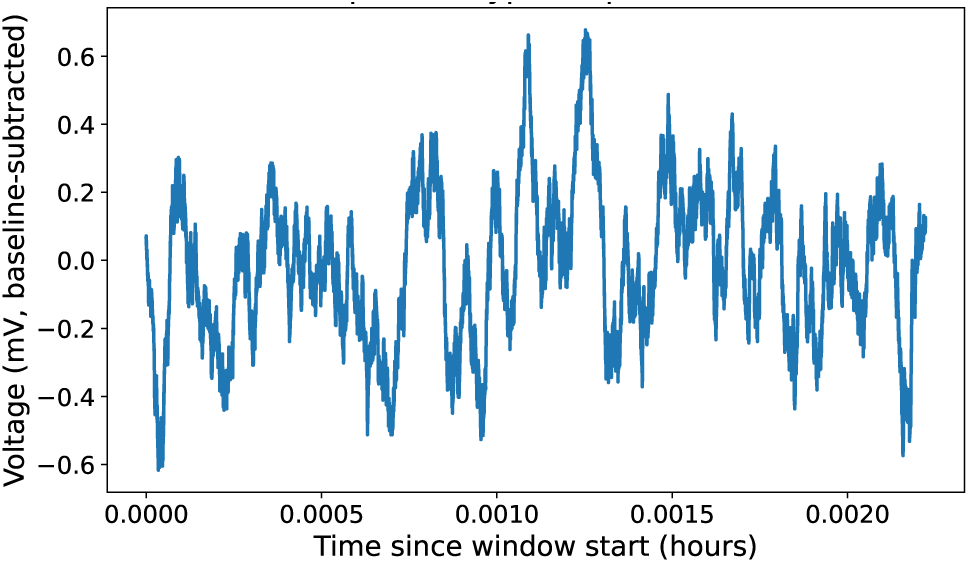
Exemplar spike train.

Across all channels, spike trains were sparse on the scale of seconds but structured on the scale of minutes to hours. ISIs spanned from ∼ 10^3^ s (tens of minutes) to ∼ 10^4^ s (several hours), indicating that spiking is dominated by slow dynamics. Pooling all channels, the median ISI was approximately 6.5 × 10^3^ s (≈ 1.8 h), the mean ISI was approximately 8.3 × 10^3^ s (≈ 2.3 h), and the standard deviation was approximately 5.8 × 10^3^ s (≈ 1.6 h). The ISI distribution was heavy-tailed (Fig. 5(a)), with occasional long silent intervals separating periods of more frequent activity.

**Figure 5:**
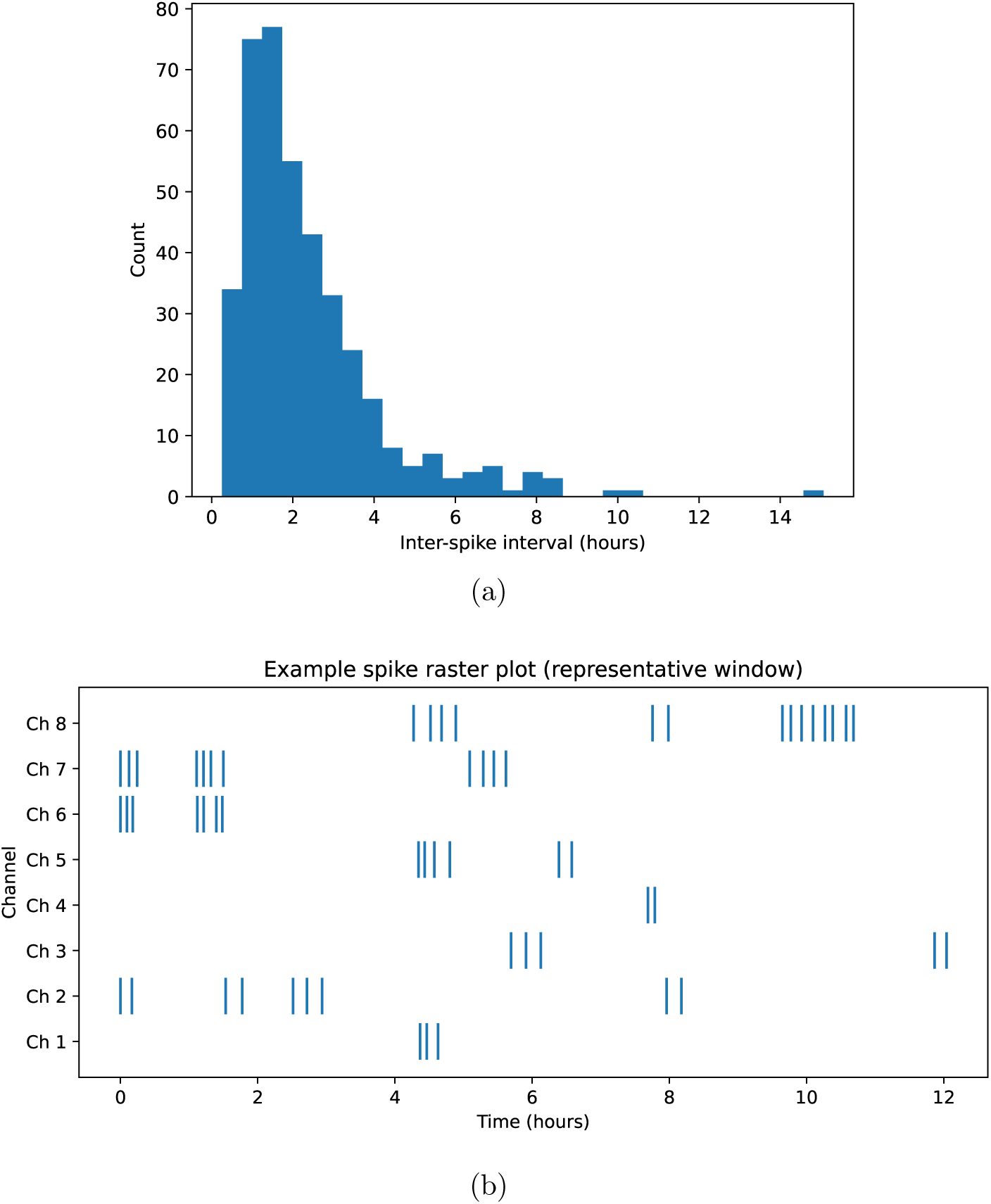
Spike-train characteristics. (a) Distribution of inter-spike intervals (ISIs) pooled across all channels, showing heavy-tailed structure on time scales of minutes to hours. (b) Example spike raster plot (onset times) for the eight channels over a representative window, illustrating intermittent clustering and long silent periods.

To quantify the regularity of spiking, we computed the coefficient of variation CV = *σ*_ISI_*/µ*_ISI_ of the ISI distribution. Across channels, CV typically exceeded unity (pooled estimate CV ≈ 0.7–1.2 depending on channel), indicating substantial variability and deviation from a near-periodic (oscillatory) process. This variability was consistent with spike generation in an excitable biological medium rather than a strictly periodic oscillator.

We next examined burst-like clustering. Bursts were defined operationally as sequences of two or more spikes separated by ISIs shorter than 30% of the channel’s median ISI (Methods). Using this criterion, spike trains showed intermittent clustering: bursts occurred but did not dominate activity. Across channels, approximately 15–35% of spikes occurred within bursts, while the remaining spikes were isolated events separated by longer ISIs. Bursts typically contained 2–4 spikes and were followed by longer recovery intervals.

For a compact visual summary, Fig. 5(b) shows representative spike rasters for all eight channels over a selected time window, illustrating (i) extended silent periods, (ii) transient episodes of increased spiking, and (iii) partial temporal alignment of events across channels. These features motivate the multi-channel propagation analysis presented in the next subsection.

### 3.3. Propagation of signal along the mycelium network

Electrical activity was recorded using a linear array of eight channels, la-belled sequentially as channels 1,2,. . . ,8. Each channel corresponds to a fixed spatial position along the mycelial network. This ordered arrangement allows direct testing of whether electrical activity propagates along the network rather than appearing independently at each recording site.

To analyse propagation, we compared spike onset times across channels. For each channel *c*, spike onsets were represented by an ordered set 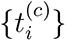. For every spike detected on channel *c*, we searched for spikes on neighbouring and non-neighbouring channels occurring within a temporal window 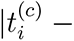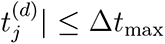, with Δ*t*_max_ = 300 s. For each associated spike pair, the delay 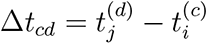 was computed.

Delay distributions were not uniform across channel pairs. Instead, delays showed clear structure that depended on the relative positions of channels in the array. In particular, spike events tended to appear first on lower-numbered channels and subsequently on higher-numbered channels, with delays increasing systematically with channel separation.

Figure 6(a) shows pooled delay distributions for adjacent channel pairs (*c, c* + 1). These distributions exhibit pronounced peaks at positive delays, indicating that spikes on channel *c* typically precede spikes on channel *c* + 1. The characteristic delays were on the order of several minutes and increased for channel pairs with larger spatial separation.

To visualise global propagation structure, we computed the median delay Δ̃*t_cd_* for each ordered channel pair (*c, d*). The resulting lead–lag matrix is shown in Fig. 6(b). The matrix reveals a clear monotonic pattern: channels with lower indices tend to lead channels with higher indices, while the opposite ordering is rare.

**Figure 6:**
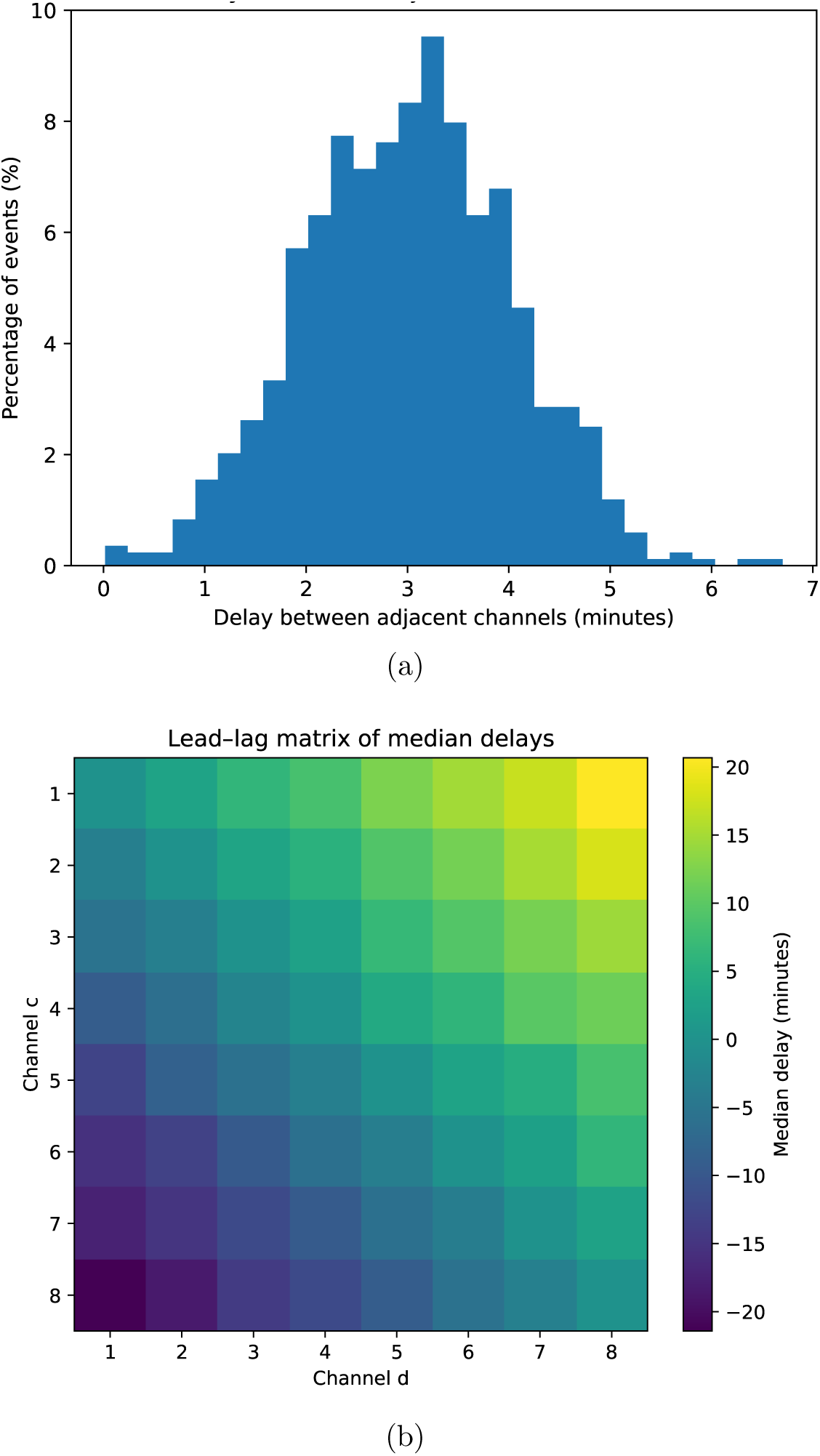
Propagation of electrical activity along the electrode array. (a) Distributions of delays between adjacent channels (c, c + 1), showing positive peaks indicative of directional propagation. (b) Lead–lag matrix of median delays between ordered channel pairs, revealing monotonic ordering from channel 1 to channel 8. (c) Median delay as a function of channel separation |d − c|, demonstrating increasing delay with distance along the array.

This monotonic lead–lag structure is incompatible with simultaneous activation or global artefacts, which would produce delay matrices symmetric around zero. Instead, the observed asymmetry indicates directional signal propagation along the channel array.

To further test the propagation hypothesis, we examined how delay magnitude depends on channel distance |*d* − *c*|. Median delays increased approximately monotonically with increasing channel separation, as shown in Fig. 7. This scaling behaviour is expected if electrical activity travels through the mycelial network at a finite speed, accumulating delay as it passes successive electrodes. Using the median delay of approximately 180 s between adjacent channels separated by approximately 2 cm, the average propagation speed of electrical activity along the mycelial network was estimated to be approximately 0.7 cm/min (approximately 40 cm/h).

**Figure 7:**
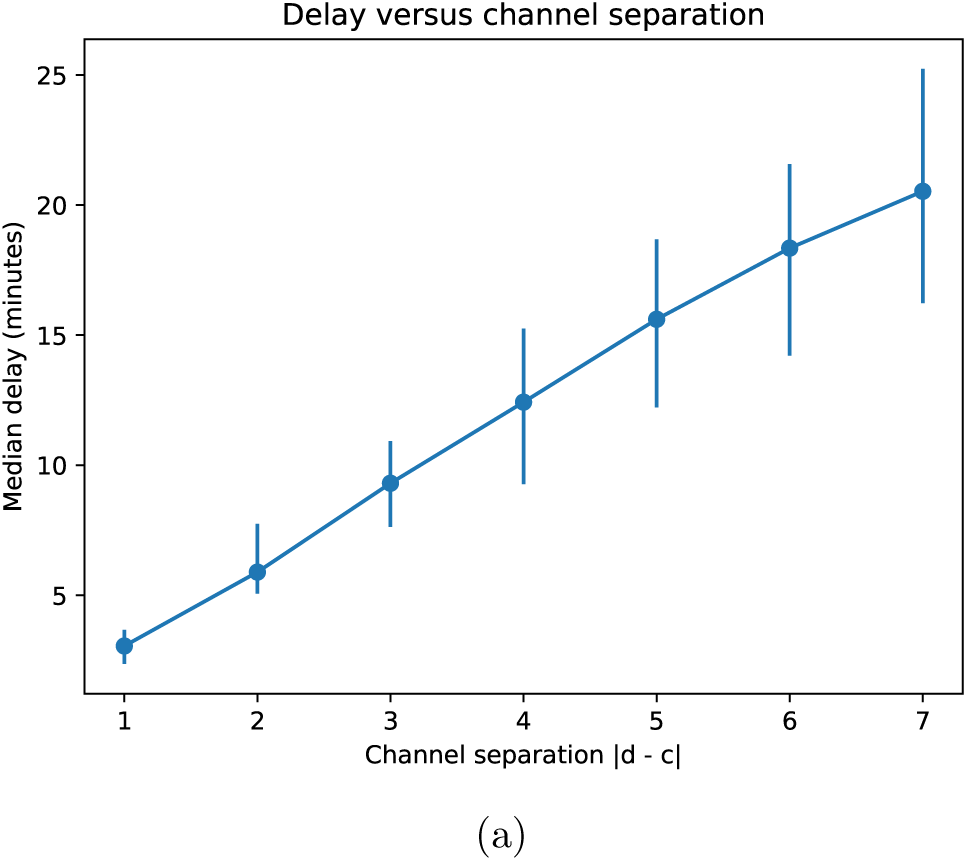
Median delay as a function of channel separation |d−c|, demonstrating increasing delay with distance along the array.

As a control, spike onset times within each channel were randomly permuted, destroying temporal relationships while preserving spike counts. After permutation, delay distributions collapsed to broad, near-symmetric profiles centred around zero, and the monotonic lead–lag structure disappeared (data not shown). This confirms that the observed propagation patterns depend on the biological timing of spikes rather than on shared drift or recording artefacts.

The results demonstrate that electrical spikes are not independent local events but propagate along the mycelial network in a directed manner detectable across the ordered electrode array. The systematic increase of delay with channel index and channel separation provides strong evidence for finite-speed signal transmission through the fungal network.

## 4. Discussion

This study provides quantitative evidence that electrical activity in substrates colonised by oyster fungi is organised in space and time and propagates along the mycelial network with a finite velocity. By combining long-term, multi-channel recordings with explicit spike detection and propagation analysis, we demonstrate that fungal electrical activity is not a collection of independent local events but forms travelling signals consistent with ionic wave dynamics.

Electrical spikes identified in the recordings are slow events with durations ranging from tens of seconds to tens of minutes and amplitudes in the millivolt range. These temporal and amplitude scales differ fundamentally from neuronal action potentials and are instead consistent with electrical phenomena reported previously in fungi, plants, and slime moulds. The asymmetric spike shape, characterised by a slow rise and longer relaxation phase, further supports an interpretation in terms of transport- and diffusion-limited processes rather than fast threshold-triggered discharges.

Analysis of spike trains reveals that electrical activity is intermittent and structured on time scales of minutes to hours. Inter-spike interval distributions are strongly right-skewed, with long silent periods separating episodes of increased activity. The high variability of inter-spike intervals, reflected in coefficients of variation close to or exceeding unity, indicates that spike generation is not periodic but governed by excitable dynamics. Burst-like clustering occurs but does not dominate activity, suggesting that the system alternates between quiescent and active states rather than operating as a sustained oscillator.

The central result of this study is the demonstration of directional propagation of electrical activity along the mycelial network. Because recordings were made using a linear array of eight spatially ordered channels, it was possible to test propagation explicitly. Delay distributions between adjacent channels show pronounced positive peaks, indicating that spikes typically appear first at one channel and subsequently at the next. Lead–lag analysis across all channel pairs reveals a monotonic ordering, with lower-numbered channels consistently leading higher-numbered channels. Importantly, delays increase systematically with channel separation, a hallmark of finite-speed signal transmission.

Using the median delay of approximately 180 s between adjacent channels separated by approximately 2 cm, the average propagation speed of electrical activity was estimated to be approximately 0.7 cm/min (approximately 40 cm/h). This velocity lies within the range expected for ionic and metabolic wave propagation in biological tissues and is comparable to values reported for electrical and calcium waves in fungal hyphae, slime mould plasmodia, and plant tissues. Such speeds are orders of magnitude slower than neuronal conduction velocities.

Control analyses based on random permutation of spike times abolish the observed delay structure and lead–lag ordering, confirming that propagation patterns arise from the biological timing of spikes rather than from shared drift, instrumental artefacts, or coincidental co-activation. Together with the long-term stability of the signals over approximately fifteen days of continuous recording, this provides strong evidence that the observed electrical activity reflects intrinsic dynamics of the living mycelial network.

The results support the interpretation of the fungal mycelium as a spatially extended excitable medium in which electrical signals propagate as ionic waves. In such a system, information is not encoded in fast all-or-none impulses but in the timing, duration, and spatial progression of slow electrical events. This view aligns with growing evidence that fungi, like plants and slime moulds, employ distributed electrical signalling to coordinate physiological processes across large spatial scales.

More broadly, the demonstrated ability of fungal networks to support propagating electrical signals strengthens the case for viewing mycelia as substrates for unconventional forms of information processing. Finite-speed wave propagation, temporal integration over long intervals, and spatially ordered signal transmission are all features that can support computation and decision-making without specialised nervous systems. Future work combining electrophysiology with imaging of ionic fluxes and detailed mapping of mycelial architecture will further clarify the mechanisms underlying these signals and their functional roles.

This study shows that electrical activity in oyster fungi is organised, propagative, and quantitatively consistent with ionic wave dynamics in a living network. These findings place fungal electrical signalling firmly within the broader class of non-neural biological information-processing systems and provide a foundation for further exploration of fungal sensing, communication, and computation.

## Acknowledgement

This project has received funding from the European Union’s Horizon 2020 research and innovation programme FET OPEN “Challenging current thinking” under grant agreement No 858132.

